# Near-critical chromatin fluctuations facilitate long-range contacts in *Drosophila* chromosomes

**DOI:** 10.1101/2025.06.20.660702

**Authors:** Gautham Ganesh, Jean-Bernard Fiche, Marcelo Nöllmann, Julien Mozziconacci

## Abstract

Recent live imaging in *Drosophila* embryonic nuclei revealed anomalous dynamics of synthetic enhancer–promoter locus pairs, challenging classical polymer models. To identify the physical mechanisms underlying this behavior, we performed coarse-grained polymer simulations exploring three chromatin organization modes: an ideal polymer, loop extrusion, and compartmental segregation. We found that genome compartments, when tuned near the coil–globule phase transition, best captured both structural and dynamical observables. Importantly, we demonstrate that the agreement with experimental data is maximized in a narrow regime characterized by near-critical polymer behavior. Incorporating loop extrusion does not markedly improve the overall fit, suggesting that the mechanism plays a limited role in shaping chromatin organization and dynamics in the *Drosophila* embryo. These results support a picture in which chromatin operates in a near-critical physical regime, enabling efficient exploration of nuclear space and frequent long-range interactions.

## Introduction

The three-dimensional (3D) organization and dynamics of chromosomes are key regulators of gene expression in eukaryotes. Chromatin is hierarchically structured within the nucleus, from megabase-scale active and inactive (A/B) compartments to sub-megabase topologically associating domains (TADs) [1–6]. These features arise through distinct molecular mechanisms.

Compartments are strongly associated with epigenetic states and transcriptional activity, yet their formation appears to involve multiple, possibly redundant, processes that lead to the segregation of different chromatin types [7, 8]. In both mammals (e.g., *Mus musculus*) and insects (e.g., *Drosophila melanogaster*), polymer models that incorporate preferential interactions between regions with similar epigenetic marks can robustly recapitulate compartmentalization patterns [9, 10].

While the mechanistic link between compartmentalization and histone modifications appears conserved in metazoans, the formation of TADs differs between species. In *Drosophila*, TADs likely emerge from the same mechanisms that drive compartmentalization [10]. In contrast, mammalian TADs are primarily shaped by loop extrusion [11–13], a process in which Structural Maintenance of Chromosomes (SMC) complexes, such as cohesin, extrude chromatin loops until halted by convergently oriented CTCF binding sites, where loops are stabilized [14–16]. Experimental and computational studies support the coexistence of loop extrusion and compartmental segregation in mammals, with loop extrusion dominating at sub-megabase scales and compartmentalization prevailing at larger genomic distances [17–19].

In contrast, the evidence for loop extrusion as a major mechanism of chromatin organization in *Drosophila* remains debated [20–22]. In flies, insulating regions are often enriched at TAD borders and can have locus-specific architectural roles [23]. Many TAD borders also coincide with transitions in chromatin state, although a subset are associated with CTCF and other insulator proteins [4, 24, 25]. Together, these observations support the view that chromatin-statedependent interactions make an important contribution to TAD formation in *Drosophila* [22].

The 3D organization of the genome plays a critical role in cell-type-specific gene regulation during development [26–28]. TADs often encompass genes promoters and their regulatory elements, such as enhancers, to facilitate their encounter while preventing unwanted contacts [29– 31]. This view of TADs as regulatory domains has helped explain how boundary disruptions can lead to ectopic gene expression [26, 32–34], though exceptions have been described [35–39]. To better understand the regulatory role of TADs, single-cell and time-resolved approaches have been instrumental. These studies suggest that TADs and compartments represent statistical rather than fixed structural units of genome organization [40], and that the internal dynamics of gene domains are key to modulating promoter–enhancer communication [41].

A recent study by Brückner and Chen et al. used live imaging in *Drosophila* embryos to simultaneously track synthetic enhancer–promoter (E–P) dynamics and transcriptional activity across genomic distances ranging from *s* = 58 kb to 3.3 Mb (Fig. 1A) [42]. These data enabled the inference of chromatin dynamics, which were then compared to predictions from polymer physics models. As expected, the motion of individual loci followed sub-diffusive behavior consistent with the Rouse polymer model, in line with previous observations in other eukaryotes [43–45]. However, the two-locus mean-squared displacement deviated markedly from predictions of standard polymer models in the field, especially for loci separated by large genomic distances [42]. These findings suggested that long-range enhancer–promoter interactions may be more prevalent than previously thought, but the underlying mechanisms remained unclear. In this study, we aim to address this gap by exploring potential physical and biological drivers of this behavior.

**Figure 1:**
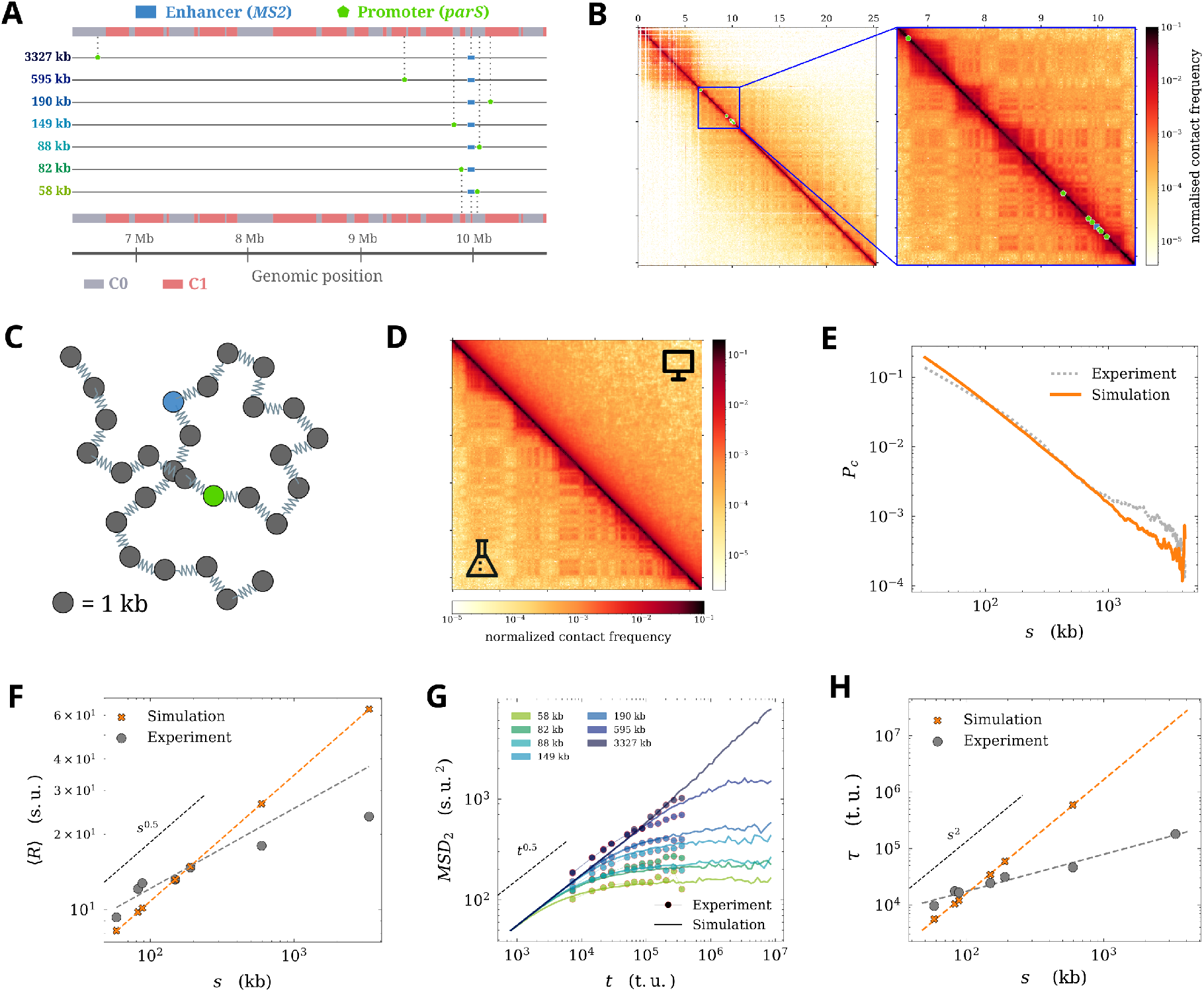
**A** – Chromosomal region modeled in this study showing insertion sites of the *MS2* (enhancer, blue rectangle) and the *parS* (promoter, green pentagon) from the seven fly lines of [42]. The compartments C0/C1 of the chromosomal region, computed from the contact map in B using eigenvector analysis (Methods), are provided with indications of which compartment the insertions fall into. **B** – Hi-C contact map of chromosome 2R (left, from [52]) with the region modeled in the present study enlarged (right). The enhancer and promoter sites use the same markers as in A. **C** – Schematic of an ideal chain, i.e., a bead-spring polymer chain without excluded volume. For illustrative purposes, the beads corresponding to the enhancer and promoter sites are marked in blue and green, respectively. **D** – Comparison between the Hi-C map from the experiment and the ideal chain simulation. **E** – Fit for P_c_(*s*) between the ideal chain simulation and the experiment. **F** – Fit for R(*s*) between the ideal chain simulation and the experiment. Theoretical behavior for an ideal chain is indicated by the black dashed line. The gray dotted line shows the linear fit to experimental data (slope *β* = 0.32) and the orange dotted line to that of the simulation (*β* = 0.50). **G** – Fit for MSD_2_(*t*) for the seven values of genomic distance *s* between the simulation and the experiment. The black dashed line shows theoretical power-law behavior. **H** – Fit for *τ* (*s*) between the simulation and the experiment. *τ* (3327 kb) cannot be computed since the corresponding MSD_2_ curve does not reach a plateau. Dashed lines show the linear fit to the data (slope *β* = 2.0 for the simulation, and *β* = 0.7 for the experiment). Black dashed line indicates theoretical power-law behavior for an ideal chain.

We modeled and simulated the dynamics of a large chromosomal region *in silico*, incorporating two well-established genome folding mechanisms—compartmental segregation and loop extrusion. We compared our simulation results to available experimental measurements, which included synthetic enhancer–promoter dynamics and physical distance, as well as Hi-C contact maps and contact probability scaling. We demonstrate that a simple heteropolymer model incorporating compartments shows good agreement with the experimental data. Notably, the best agreement between simulations and experiments occurs near the coil–globule phase transition, where the chromatin polymer fluctuates between extended and compact configurations [46, 47]. These findings suggest that compartmental segregation operating near this transition provides a mechanistic basis for the observed structure and dynamics of *Drosophila* chromosomes.

## Results

We aimed to investigate the mechanisms underlying the structural and dynamical properties of chromosomal loci reported by Brückner et al. [42]. Since polymer simulations have been widely used alongside experiments for this purpose [48–51], we simulated a large region of chromosome 2R in *Drosophila* (Fig. 1A) using models that are widely used for chromatin. Specifically, we explored three models: an ideal chain, a loop-extruded polymer, and a block copolymer.

We fit simulation outputs to corresponding experiments using three key observables: (i) the probability of contact P_c_(*s*) as a function of genomic separation *s* from Hi-C data acquired in the same developmental stage (nuclear cycle 14) [52] (Fig. 1B), (ii) the mean E–P physical distance R(*s*), and (iii) the two-locus mean-squared displacement MSD_2_(*t*) for varying *s* [42] (see Methods). The first two observables capture structural properties, while the third describes chromatin dynamics. After fitting, we also included other observables from Brückner et al., such as the single-locus mean-squared displacement (MSD_1_(*t*)) and the relaxation time (*τ* (*s*)) (see Methods) for visual comparison of best fits with the experimental results. We note that the experimental assay probes an engineered regulatory system rather than a simple endogenous one-to-one enhancer–promoter interaction, and its interpretation therefore requires accounting for the additional *homie* elements. All observables were thus computed only from experimental trajectories exhibiting an open (i.e. with no loop) configuration (denoted the O_off_ state in Brückner et al.) to exclude persistent contacts due to the presence of *homie* loop-forming elements [53] in the experimental setup.

To evaluate model performance, we quantified deviations between simulations and experiments using three metrics: 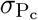 for P_c_(*s*), *σ*_R_ for R(*s*), and 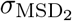 for MSD_2_(*t*) (see Methods). We scaled both spatial and temporal units to match the experimental data for the three observables and computed the deviations *σ* between the simulation and the experiment (see Methods). For each model, we identified parameter sets minimizing these deviations, compared deviation values, and visually assessed the fits to determine consistency with the experimental data.

### An ideal chain poorly fits the experimental data

We first applied our approach to the simulation of an ideal chain, a basic polymer model with well-established theoretical predictions. We modeled the 4.2 Mb segment of chromosome 2R encompassing all synthetic enhancer-promoter pairs as a bead-spring polymer without excluded volume (Fig. 1C) and simulated it as described in Methods.

The simulated Hi-C map, computed using a contact threshold of *k* = 250 nm (see Methods), showed a decay in contact frequency with increasing genomic distance *s*, but lacked the structured patterns observed in the experimental map (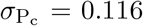 ; map-map Pearson correlation *r* = 0.01 Fig. 1D,E). The observed scaling exponents for R(*s*) and MSD_2_(*t*) were both close to 0.5 (Fig. 1F,G), consistent with ideal chain behavior [54]. The mean spatial distance R(*s*) deviated from the experimental trend (*σ*_R_ = 0.848; Fig. 1F), and while the fit for MSD_2_(*t*) was reasonable for genomic separations up to 190 kb, it diverged at larger distances (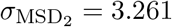 ; Fig. 1G), highlighting the limitations of the ideal chain in capturing long-range chromatin behavior. The relaxation time *τ* (*s*), which indicates the crossover time between the initial growth regime and the plateau of MSD_2_(*t*) (see Methods), provides an additional measure of how rapidly pairs of loci explore their available spatial range. For an ideal chain, theory predicts a steep scaling with genomic distance, *τ* (*s*) ∼ *s*^2^, whereas the experimental data show a much weaker dependence with *τ* (*s*) ∼ *s*^0.7*±*0.2^, indicating a clear disagreement between the model and the observed behavior (Fig. 1H).

To test whether excluded volume effects could improve the model, we added steric interactions to the ideal chain. However, this modification further increased the discrepancies with the experimental data (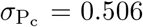 , *σ*_R_ = 1.086, 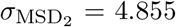), as also confirmed by visual inspection of the fits (Fig. S1A–C). These results indicate that the ideal chain, with or without excluded volume, fails to capture the structural and dynamic features observed, motivating the exploration of more complex models.

### Loop extrusion marginally improves the fit to data

Extensive simulation-based and experimental studies in mammals have shown that loop extrusion by chromatin-associated factors, such as cohesins, affects both chromatin compaction and its diffusive scaling behavior [55–57]. We hypothesized that loop extrusion could promote globular chromatin organization while preserving near-ideal chain dynamics. To test this, we implemented loop extrusion using a well-established two-step framework [13, 58]: first, the positions of loop-extruding factors (LEFs) were determined on a one-dimensional (1D) lattice, and then these loop constraints were applied to a 3D bead-spring polymer simulation (Fig. 2A; see Methods).

**Figure 2:**
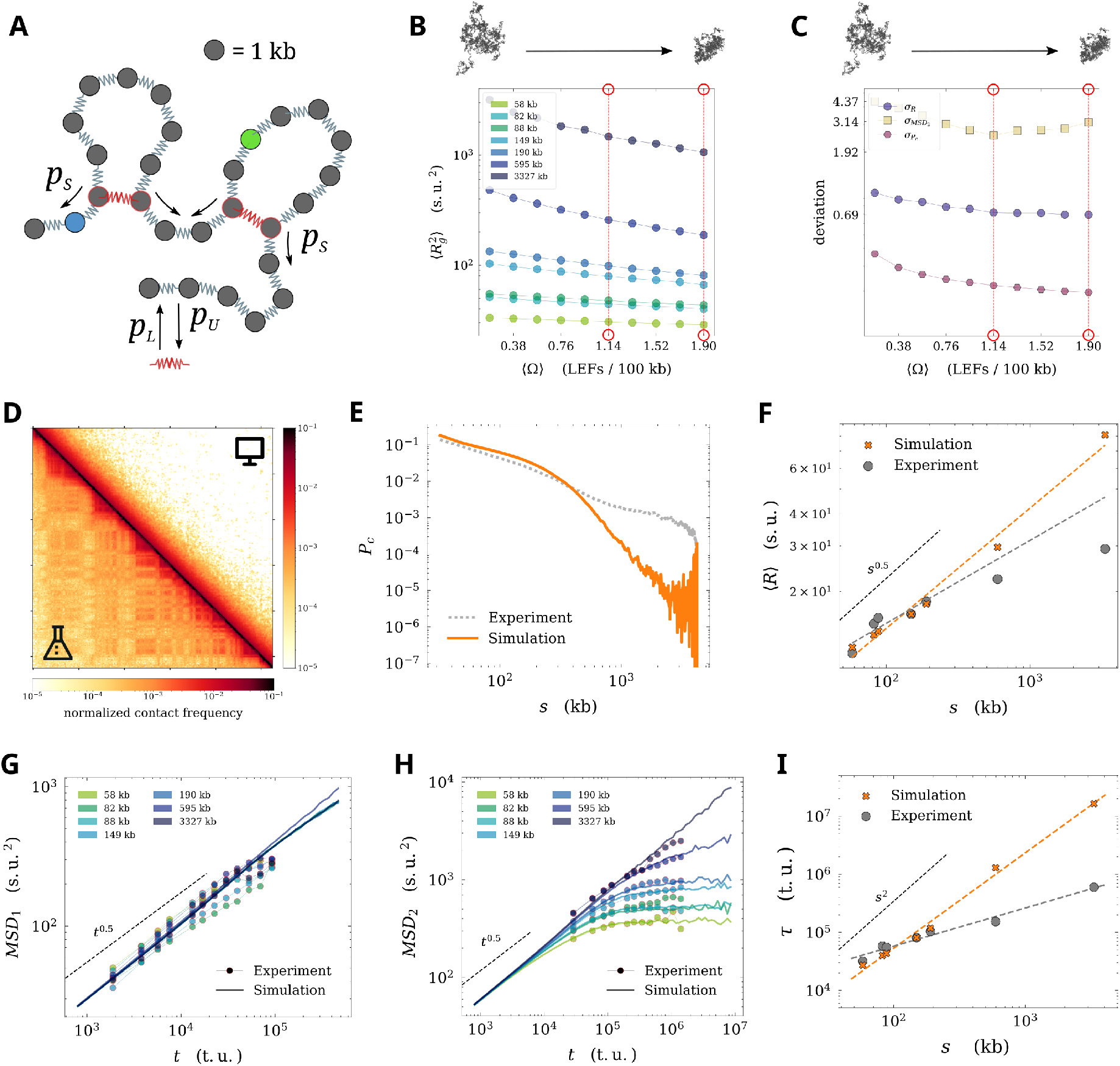
**A** – Positions of LEFs from the lattice-based stochastic simulation (springs marked in red) are integrated into the bead-spring polymer chain with excluded volume (Methods). LEF activity is controlled by the parameters *p*_*U*_ , *p*_*L*_, and *p*_*S*_, which correspond to LEF unloading, loading, and stepping probabilities, respectively. **B** – Mean-squared radius of gyration 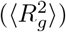 for each enhancer-promoter segment under increasing mean LEF occupancy ⟨Ω⟩ (in numbers per 100 kb). Red circles and vertical dotted lines indicate the ⟨Ω⟩ at which the best fits to experimental data occur (see panel C). **C** – Deviations between experiment and simulation for R(*s*), MSD_2_(*t*) and P_c_(*s*) as a function of increasing ⟨Ω⟩. Minima marked with red circles and dotted lines. **D** – Comparison of Hi-C maps between experiment and best-fitting simulation for Hi-C. **E** – Best fit of P_c_(*s*) between simulation and experiment. **F** – Best fit of R(*s*) between simulation and experiment. The black dashed line indicates ideal chain behavior. Orange and gray dotted lines show linear fits to simulated and experimental data, respectively (*β* = 0.47 for the simulation). **G** – Best fit of MSD_1_(*t*) between simulation and experiment. The black dashed line shows the ideal chain exponent. **H** – Best fit of MSD_2_(*t*) between simulation and experiment. The black dashed line shows the ideal chain exponent. **I** – Best fit of *τ* (*s*) between simulation and experiment. Dashed and dotted lines same as described before. The scaling exponent from the simulation *β* = 1.67.

We explored a range of LEF occupancies (⟨Ω⟩) from 0.19 to 1.9 LEFs per 100 kb, consistent with previous estimates [13, 59]. LEF dynamics were governed by four parameters (Fig. 2A): the loading probability *p*_*L*_ (varied to control ⟨Ω⟩), unloading probability *p*_*U*_ = 0.01, sliding probability *p*_*S*_ = 0.5, and crossing probability *p*_*C*_ = 0.5, which allows LEFs to bypass one another along the chromatin fiber (see Methods for details). By fixing *p*_*U*_ and *p*_*S*_, we maintained a mean loop size of approximately 100 kb, close to in vivo estimates in other organisms [45].

Although TAD boundaries in *Drosophila* are often associated with insulator proteins [21], whether these elements act as barriers to loop-extruding factors in vivo remains unclear. In the absence of a well-established blocking rule for *Drosophila*, we did not impose boundary constraints in our simulations and instead treated LEFs as freely moving with the possibility of crossing as a simplifying assumption.

To monitor polymer compaction across simulations with varying LEF occupancy, we measured the mean-squared radius of gyration 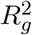 (see Methods) for all enhancer–promoter segments (Fig. 2B). As expected, 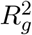 decreased with increasing mean LEF occupancy, indicating progressive chromatin compaction. We then evaluated how each model’s agreement with experimental data changed as LEF occupancy increased by tracking the three deviation metrics between simulations and experiments (Fig. 2C). The structural deviation *σ*_R_ decreased with increasing ⟨Ω⟩ and reached a minimum at 1.9 LEFs per 100 kb (*σ*_R_ = 0.692). In contrast, the dynamic deviation was minimized at an intermediate occupancy of ⟨Ω⟩ = 1.14 LEFs per 100 kb with 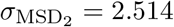, marking an improvement over the ideal chain. The Hi-C deviation 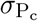 remained relatively constant across the tested range. These results suggest that although loop extrusion may contribute to chromatin compaction and can partially reconcile structural and dynamic features observed in vivo, no single occupancy value optimally fits all observables.

The Hi-C map obtained from the simulation with the lowest discrepancy lacks the long-range contacts observed in the experimental data (Fig. 2D, Pearson *r* = −0.02). Consistently, the contact probability P_c_(*s*) is not well captured at large genomic distances, where the simulated curves decay too rapidly compared to the experimental trend (Fig. 2E). The agreement for R(*s*) is improved relative to the ideal chain, and the experimental scaling exponent is reproduced up to *s* = 190 kb. However, R(*s*) increases markedly at larger genomic distances, reflecting the limited influence of loop extrusion at these scales in our simulations (Fig. 2F).

The single-locus dynamics MSD_1_(*t*) remain close to ideal polymer behavior (Fig. 2G), indicating that loop extrusion has only a modest impact on local dynamics. A similar behavior is observed for MSD_2_(*t*), where the model accurately reproduces the curves for *s* = 58 to 190 kb, but fails to capture the rapid plateauing observed experimentally at larger separations (*s* = 595 and *s* = 3327 kb; Fig. 2H), highlighting incomplete agreement with long-range dynamics despite the improvement in 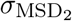. Finally, the relaxation time *τ* (*s*) exhibits a much steeper scaling with genomic distance than observed experimentally (Fig. 2I), further emphasizing the limitations of loop extrusion alone in reproducing chromatin dynamics.

After the fitting procedure, we extracted the scaling factors to convert simulation spatial and temporal units (s.u. and t.u.) into experimental units (nm and s) based on the best fit to MSD_2_(*t*). Using these, we estimated that the mean LEF sliding speed was approximately 0.8 kb/s, consistent with in vitro measurements of cohesin velocity [60]. The LEF unloading probability *p*_*U*_ corresponded to a mean residence time of ∼ 63 s, substantially shorter than experimental estimates of ∼ 360 s obtained under CTCF-depleted conditions in mouse embryonic stem cells [45, 61].

Taken together, our simulations indicated that loop extrusion contributes to chromatin compaction and improves agreement with some observables, yet it is insufficient on its own to fully explain the structural and dynamical features observed in the experimental data.

### Chromatin structure and dynamics are consistent with compartmental segregation

Next, we investigated the effect of compartmental segregation on chromatin structure and dynamics. To identify chromatin compartments, we extracted the first eigenvector of the experimental Hi-C map at 16 kb resolution after normalization by genomic separation (Fig. 1B), assigning each genomic bin to one of two compartments, C0 or C1 (see Methods). This analysis revealed alternating compartment blocks of varying genomic lengths within our region of interest (Fig. 1A, bar marked with red and gray regions). We then assigned each bead in our polymer model to compartment C0 or C1 based on its corresponding genomic bin (Fig. 3A, left). To model compartmental interactions, we introduced a distance-dependent attractive potential between beads belonging to the same compartment, following established block copolymer approaches [10, 49] (Fig. 3A, right). For simplicity, we assumed a uniform interaction energy *ε* for both C0–C0 and C1–C1 bead pairs.

**Figure 3:**
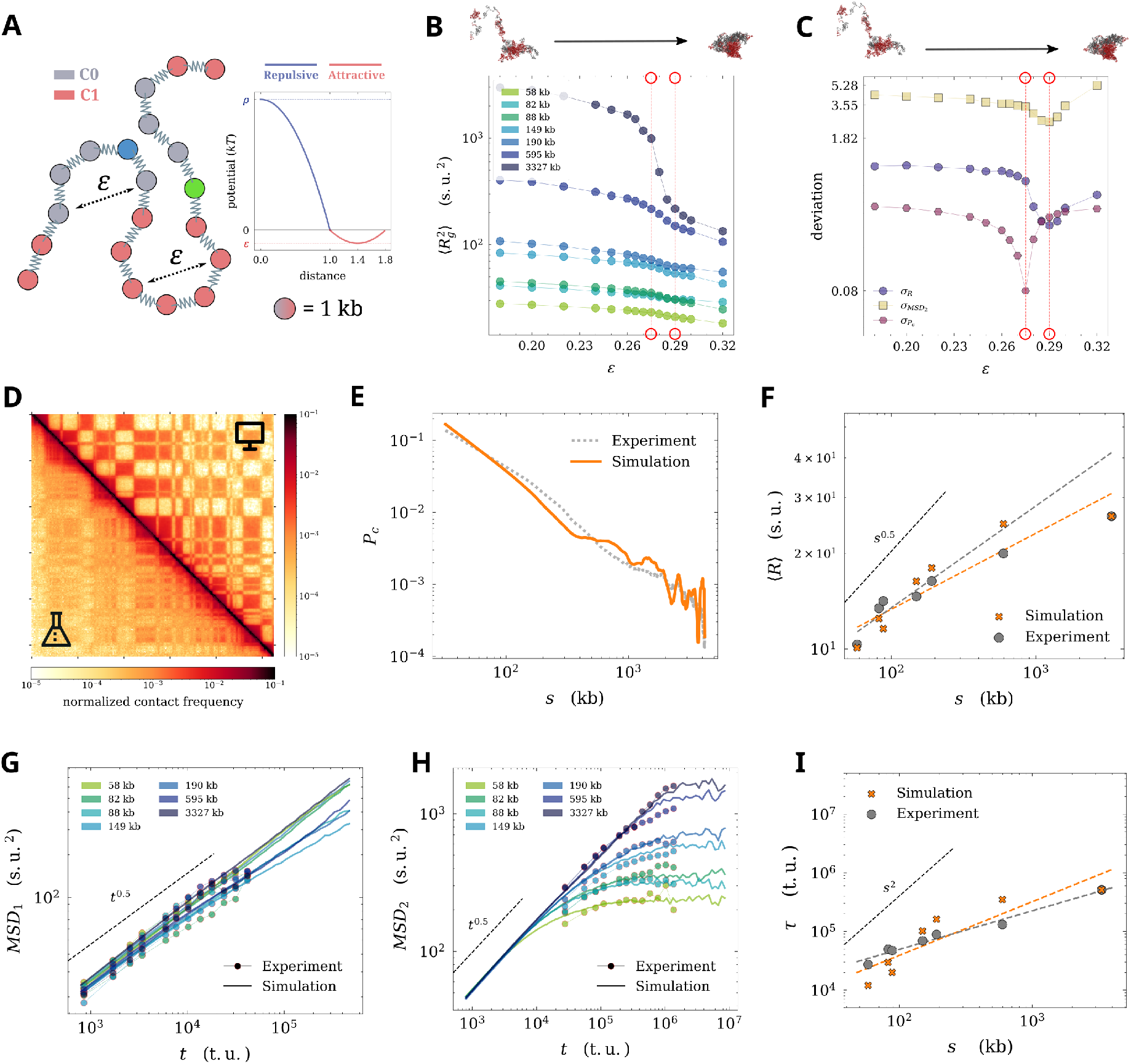
**A** – (Left) Bead-spring block copolymer chain with excluded volume and two bead types colored based on the compartment in which they belong. The beads corresponding to the enhancer and promoter sites are marked as before. (Right) The short-range attractive potential between beads of the same type is given by the parabolic well of depth *ε* (in red), while the repulsive interactions enforce excluded volume (in blue) for all beads. Interactions between beads of different types are only repulsive. **B** – Radius of gyration of each enhancer-promoter segment as a function of compartmentalization strength *ε*. Values of *ε* at which the best fits with experimental data occur (from the following panel) are indicated as in 2. **C** – 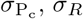, and 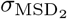 as a function of *ε*. The positions of the occurrence of best fits are marked in red. **D** – comparison of Hi-C contact maps between simulation and experiment for the best fit to Hi-C. **E** – Best fit of P_c_(*s*) between simulation and experiment. **F** – Best fit with experimental R(*s*) with dotted lines indicating linear fits (*β* = 0.24 for the simulated data). The black dashed line shows ideal chain exponent. **G** – Best fit of MSD_1_(*t*) between simulation and experiment. The black dashed line shows the ideal chain exponent. **H** – Best fit with experimental MSD_2_(*t*). The black dashed line indicates ideal-chain behavior, as above. **I** – Relaxation time computed from the best-fit simulation to R(*s*) and MSD_2_(*t*). Dashed lines indicate scaling exponents, *τ* (*s*) ∼ *s*^0.91^ for the simulation.

While systematically varying the strength of the attractive potential *ε* in our simulations, we observed that each enhancer-promoter polymer segment underwent a characteristic coil–globule transition (i.e., larger to smaller mean 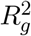, Fig. 3B) [46], consistent with previous studies of self-attracting polymers [47, 62]. To identify the value(s) of *ε* that best matched the experimental data, we computed the deviations *σ*_R_, 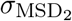, and 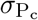 across a range of *ε* values. Both *σ*_R_ and 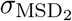 reached minima at *ε* = 0.29, while 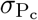 was minimized at a slightly lower value of *ε* = 0.275 (Fig. 3C). Overall, this model yielded significantly better overall agreement with experimental data than previous models 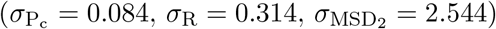.

The Hi-C map displayed the checkerboard pattern characteristic of compartments, resulting in increased correlation with the experimental data despite higher contrast in the simulated map (Fig. 3D, Pearson *r* = 0.43). The contact probability P_c_(*s*) captured both short- and long-range structural features of the polymer; however, while the model reproduces the overall trend of the experimental data, deviations remain, in particular at genomic distances between 100 kb and 1 Mb, where the simulated decay is steeper than observed (Fig. 3E). The R(*s*) curve from simulations follows the experimental trend across all scales (Fig. 3F).

The single-locus dynamics MSD_1_(*t*) remain close to ideal polymer behavior, fitting the experimental data well (Fig. 3G). The fit to MSD_2_(*t*) shows notable improvement over the Rouse model; importantly, in contrast to previous models, the curves for *s* = 595 and *s* = 3327 kb correctly transition into the plateau, although discrepancies remain at shorter time scales for smaller *s* values (Fig. 3H). For the best-fit simulation to dynamics (i.e., the one minimizing 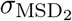), we measured a scaling exponent *τ* (*s*) ∼ *s*^0.91^ for the relaxation time, much closer to the experimental value of 0.7 ± 0.2 than the ideal-chain prediction, and therefore consistent with the observed weak dependence of relaxation time on genomic distance (Fig. 3I) [42].

Collectively, these results indicate that compartmental segregation accounts for chromatin structure, dynamics, and relaxation reasonably well and reproduces them more accurately than either the ideal-chain or loop-extrusion models, suggesting that it is a major determinant of megabase-scale chromatin organization and dynamics at this stage of embryonic development.

### Evidence for near-critical chromatin dynamics

The best agreement between simulations and experimental data was obtained for values of compartment interaction strength *ε* close to the coil–globule transition (Fig. 3). Previous studies have already indicated the significance of this transition, connecting it to behaviors characteristic of critical interacting polymers, such as increased conformational fluctuations [46, 47]. To investigate whether the range of best fits reflected a specific physical regime, we systematically analyzed how two physical quantities (fluctuations of R(*s*) and the power spectral density of chromatin motion) depend on *ε*.

First, we characterized the global conformational fluctuations of polymers from our simulations by measuring the normalized standard deviation of the squared radius of gyration as a function of *ε*. This quantity remains constant until *ε* ≈ 0.24, with a pronounced peak at *ε* = 0.28, and a rapid decrease for larger *ε*, indicative of enhanced fluctuations associated with the coil–globule transition (Fig. 4A). A similar, albeit weaker, increase is observed for representative compartment domains considered independently (Fig. S2A). This result corroborates that near the transition, large conformational fluctuations are observed even at compartment domain length scales, because the polymer alternates between extended and compact states [47].

**Figure 4:**
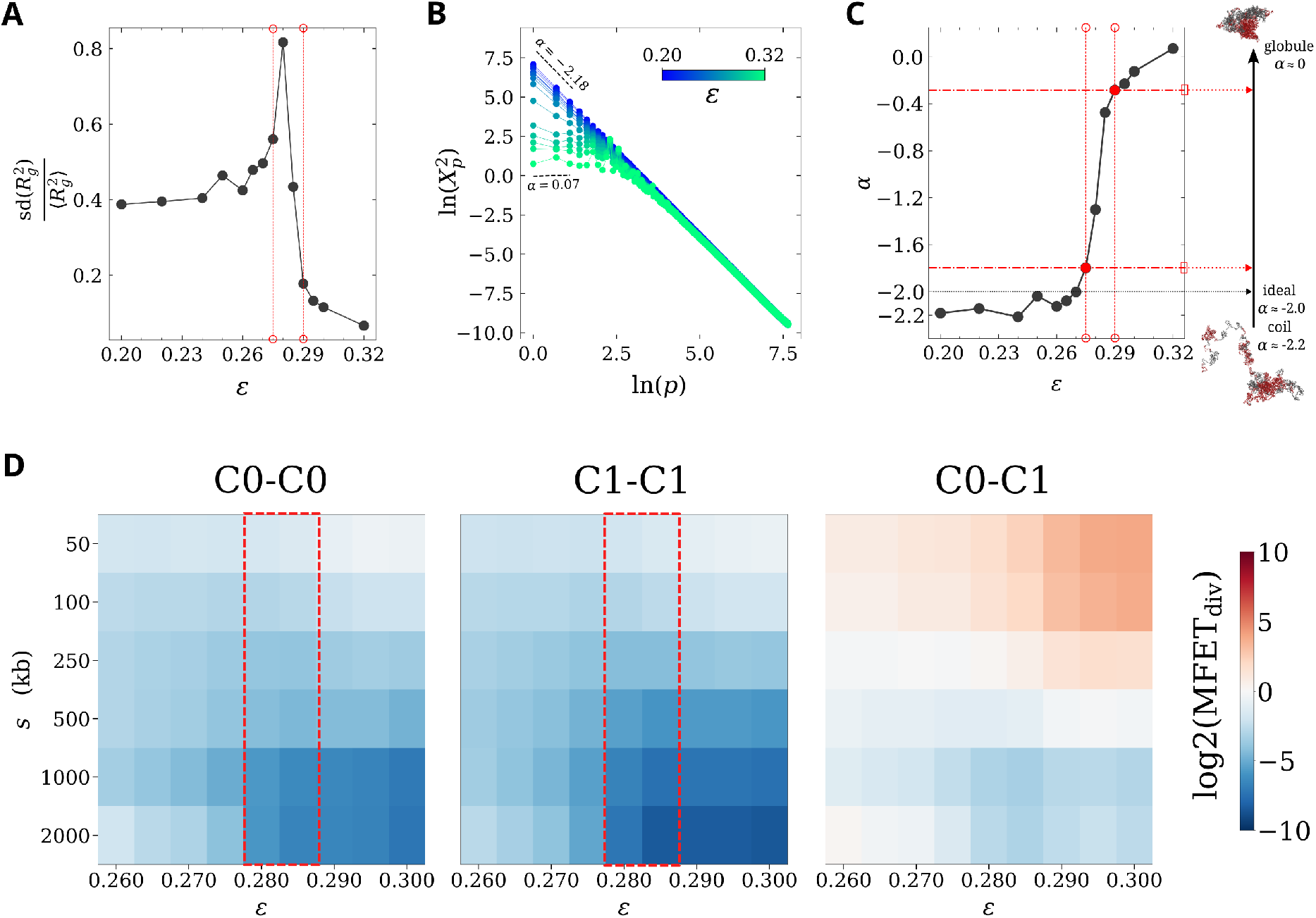
**A** – Normalized standard deviation of the squared radius of gyration 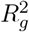 of the full chromatin block copolymer as a function of *ε*. Dotted vertical red lines indicate the best-fitting *ε* values for the compartment-based block copolymer. **B** – Power spectral density 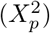 curves for polymer modes *p* computed from an ensemble of polymer conformations for increasing values of *ε*. The slope of the 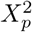 for the first three modes, called *α*, indicates the conformational ensemble’s position within the coil–globule transition. **C** – Scaling exponent of the power spectral density (*α*) for the same polymer. Dotted vertical red lines same as in A, and horizontal dashed lines indicate corresponding *α* values. The scheme on the right shows the expected *α* range from a coil to a globule. **D** – Deviations in mean first encounter times between pairs of loci from different combinations of compartment types, relative to a Rouse polymer with excluded volume for increasing values of *ε*. Regimes with increased encounter kinetics marked as dotted rectangles in C0–C0 and C1–C1 grids. Crossover between the two regimes at the coil-globule transition marked in red.

A complementary signature of this process is provided by the power spectral density (PSD) of chromatin motion [63]. Low spectral modes capture large-scale polymer structural properties. They are stronger in coil-like conformations, which therefore exhibit a steeper spectrum, and weaker in globule-like conformations, where confinement flattens the spectrum (Fig. 4B). The spectral exponent *α* therefore provides an effective indicator of the polymer state and undergoes a crossover between coil-like (*α* = -2.2) and globule-like regimes (*α* = 0) [63] (Fig. 4C). For our best-fit simulation cases of *ε* = 0.275 and *ε* = 0.290, *α* ≈ −1.8 and *α* ≈ −0.3, respectively, falling, as expected, close to the edges of the coil–globule transition. However, *α* values for the largest C0 and C1 domains occur close to -1.6 and -1.1, well within the coil–globule transition. The same transition is apparent when considering the spectral exponent for the two largest compartment domains (Fig. S2B). In fact, *α* values for compartment domains exhibit a domainsize-dependent effect—they appear closer to a globular state for larger compartmental domains, and further within the transition for smaller domains, since the coil–globule transition of finite-size polymers displays a broadened crossover rather than a sharp transition (Fig. S2C, [46]).

Together, these findings show that the chromatin properties reproduced by the model occur within a narrow parameter range close to the coil–globule transition and are consistent with near-critical polymer behavior at the scale of individual compartment domains.

### Attractive compartmental interactions facilitate homotypic encounters

To quantify more directly how compartmental interactions affect communication between distant loci, we measured mean first encounter times (MFETs) between several pairs of loci, and quantified their deviation from corresponding measurements for a reference homopolymer with no attractive interactions. Fig. 4D shows the log ratio between these two quantities as a function of genomic distance *s* and interaction strength *ε*, separately for the different combinations of compartment identities. Negative values (blue) indicate that encounters occur faster than in the reference case, whereas positive values (red) indicate slower encounter kinetics. A clear asymmetry emerges between homotypic and heterotypic pairs. For both C0–C0 and C1–C1 pairs, increasing attractive interaction strength systematically reduces the MFET, consistent with compartmentalization facilitating encounters between loci within the same compartment type (Fig. 4D, left and center grids). Conversely, heterotypic C0–C1 pairs appear to show a more complex genomic-distance-dependent effect. MFETs are increased relative to the reference case for *s* up to 500 kb, consistent with the observed segregation. For *s* greater than 500 kb, increasing *ε* reduces MFETs, as observed for homotypic encounters (Fig. 4D, right).

Our analysis also reveals two distinct regimes for homotypic encounters (Fig. 4D).In the coil state (*ϵ* ≤ 0.28), attractive interactions moderately reduce MFETs for all distances between pairs of loci. In the globule state (*ϵ* ≥ 0.29), this reduction becomes less pronounced for short distance pairs and more important for long distance pairs (*s* ≥ 500 kb). The crossover between these two regimes occurs near the coil–globule transition (i.e., *ε* ≈ 0.28 − 0.29), where encounter kinetics show reduced dependence on *s* (Fig. 4D, pink rectangles). In this regime, homotypic encounters are enhanced both at short and long genomic distances. This behavior suggests that proximity to the transition provides an efficient compromise between local exploration of chromosomal loci and global compaction, thereby facilitating long-range communication without sacrificing short-range encounter kinetics.

### Combining compartmental segregation and loop extrusion models

Compartmental segregation emerged as the main driver of chromatin structure and dynamics in our system of interest. Yet, residual discrepancies at short genomic and temporal scales suggested an additional contribution from other processes. Since loop extrusion and compartments can interact both synergistically and competitively in mammalian systems [18, 19, 64], and because our loop-extrusion simulations improved R(*s*) and MSD_2_(*t*) around ∼100 kb, we combined both mechanisms in a unified bead-spring polymer model with excluded volume (Fig. 5A). To identify optimal parameter combinations, we performed a grid search over a restricted range of compartment interaction strengths (0.275 ≤ *ε* ≤ 0.300) and mean LEF occupancies (0.19 ≤ ⟨Ω⟩ ≤ 0.95 LEFs per 100 kb), and quantified for each parameter pair the discrepancies between simulations and experiments for all three observables (Fig. S3A). We found that the addition of loop extrusion resulted in a near-universal improvement to the fits for P_c_(*s*) (Fig. S3A, left) and MSD_2_(*t*) (Fig. S3A, right) across all *ε*, whereas the fit to R only improved when both *ε* and ⟨Ω⟩ increased (Fig. S3A, center).

**Figure 5:**
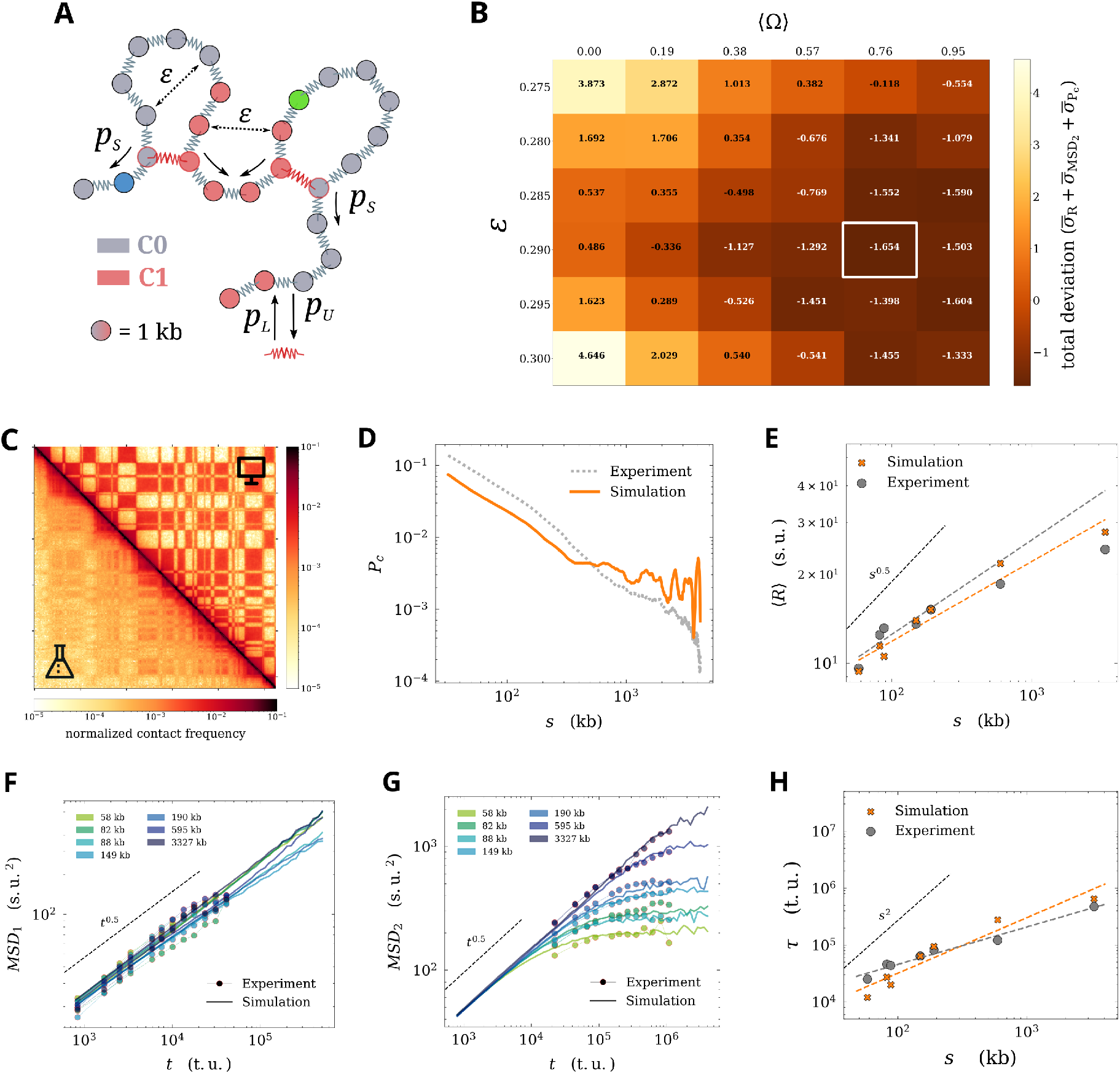
**A** – Schematic of loop extrusion on a compartment-based block copolymer. *ε, p*_*L*_, *p*_*U*_ , and *p*_*S*_ are the same as indicated previously. **B** – Grid with the sum of normalized deviations 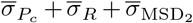 and global minimum highlighted. **C** – Hi-C map comparison between simulation and experiment for the global minimum. **D** – Best fit for P_c_(*s*) between simulation and experiment for the global minimum. **E** – fit with experimental R(*s*) for the same case; the scaling exponent from simulations is *β* = 0.27. Dashed and dotted lines are the same as indicated previously. **F** – Best fit with experimental MSD_1_(*t*). **G** – Best fit with experimental MSD_2_(*t*). **H** – Best fit with experimental *τ* (*s*); the simulated scaling exponent is *β* = 0.98.

The fit to *P*_*c*_(*s*) favored higher LEF occupancy and slightly increased compartment interaction strength relative to the best-fit compartment-only model, with a minimum at *ε* = 0.280 and ⟨Ω⟩ = 0.95 LEFs/100 kb (Fig. S3A, left). In contrast, the best fit for R(*s*) occurred at *ε* = 0.300 and ⟨Ω⟩ = 0.76 LEFs/100 kb (Fig. S3A, center). The dynamic observable MSD_2_(*t*) followed a similar trend, with a minimum at *ε* = 0.290 and ⟨Ω⟩ = 0.76 LEFs/100 kb (Fig. S3A, right grid). Each deviation metric showed improved minima 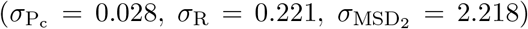 compared to the compartment-only model, and the respective visual comparison of the fits of each observable reflected this marked improvement (Fig. S3B–D, plots marked in red).

Because the optimal parameter combination differed among observables, we Z-score normalized the three deviations and summed them to identify a global optimum (see Methods). This procedure yielded a minimum at *ε* = 0.290 and ⟨Ω⟩ = 0.76 LEFs per 100 kb (Fig. 5B), corresponding to approximately one extruder per 130 kb.

Visualizing the best fits for each observable at the global optimum, we find that the Hi-C map still displayed stronger compartmentalization than observed experimentally and did not reproduce the overall contact pattern better than the previous models (Fig. 5C, Pearson *r* = 0.4). The contact probability P_c_(*s*) was likewise worse than the compartment-only model 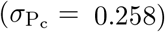, with much higher contact probabilities at the Mb scale than observed experimentally (Fig. 5D). The agreement for R(*s*) improved slightly, as reflected by the reduced deviation *σ*_R_ = 0.283 (Fig. 5E). The combined model reproduced the overall trend of the experimental data reasonably well across genomic scales, although visual inspection revealed larger deviations than in the compartment-only model at the largest genomic separations. The single-locus dynamics MSD_1_(*t*) remained consistent with the near-Rouse behavior observed experimentally (Fig. 5F). The agreement for MSD_2_(*t*) improved 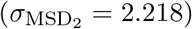, especially at short times and short genomic separations. In addition, the crossover of *s* = 595 kb and *s* = 3.3 Mb curves into the plateau was captured better than the compartment-only model (Fig. 5G). Surprisingly, despite the improvement to MSD_2_(*t*), the relaxation time scaling *τ* ∼ *s*^0.98^ did not improve compared to the compartment-only model (Fig. 5H).

Taken together, these results indicate that models combining compartmental segregation and loop extrusion can reproduce individual observables from the experiment with excellent agreement, but the global best-fit model does not improve consistency over the compartment-only model and fails to account for all experimental observations simultaneously (Fig. S4).

## Discussion

Here, we leveraged coarse-grained polymer simulations to test two well-known mechanistic models of chromatin organization and explain structural and dynamical properties of a *Drosophila* chromosome during early embryo-genesis. First, we confirmed that the ideal chain model, with or without excluded volume, cannot reproduce experimental observations. We then tested a mechanistic loop extrusion model by systematically varying the mean occupancy of loop-extruding factors while maintaining their processivity. Although this model slightly improved agreement with specific experimental observables, it failed to capture their behavior within the parameter space explored. Next, we simulated compartmental segregation using a simple block copolymer model, which reproduced the experimental data with reasonable agreement. Although discrepancies in describing the Hi-C data remain, we conclude that our results, alongside other work [4, 10, 25], are consistent with compartments playing a central role in shaping chromatin structure and dynamics.

The best agreement with experimental data occurred near the coil–globule phase transition, a regime known to display enhanced conformational fluctuations while retaining fast polymer dynamics. [46, 47]. Using measurements of polymer-size fluctuations, power spectral density, and locus encounter kinetics, we show that the experimental observations are consistent with the physical properties of a polymer operating near the coil–globule transition. Recently, such a near-critical polymer state was also shown based on analysis of imaging data both in *Drosophila* [46] and mammalian chromosomes [47]. We propose that this near-critical regime facilitates long-range interactions by reducing their dependence on genomic distance and enabling rapid spatial exploration within the nucleus. Our model thus predicts that modest perturbations of compartmental interaction strength—through epigenetic modifications, for example—lead to changes in chromatin dynamics and long-range interaction frequencies.

Lastly, our combined best-fit model of compartmental segregation and loop extrusion showed varying changes in agreement with different experimental observables, with no consistent overall improvement over the best compartment-only model in reproducing all experimental results, reiterating the important role of compartments in genome organization across kb and Mb scales. The role of loop extrusion in *Drosophila* remains debated [21, 22]; however, our work, together with others [20], even if loop extrusion occurs in flies, its contribution to the structural and dynamical observables examined here is limited.

From a polymer physics perspective, recent work has sought to reconcile observations of chromosome structure and dynamics (and even mechanics) using diverse modeling approaches [57, 65, 66]. Kadam et al. [57] used the mechanistic loop extrusion model to simulate a chromosomal locus in mouse embryonic stem cells and demonstrated that it simultaneously reproduces both Hi-C and live imaging dynamics data, presenting parallel work similar to the approach used in our study but in a mammalian system. Shi et al. [66] used a maximum-entropy model to generate and analyze structural ensembles from a single Hi-C contact frequency map; this approach, while not mechanistic, could reproduce observations from Brückner et al. Our work goes beyond this effort by offering a biologically grounded, mechanistic interpretation of *Drosophila* chromatin organization based on a well-established physical modeling approach, paving the way for the generation of testable predictions for future experiments.

We focused here on two classes of mechanisms that have been shown to robustly account for genome folding across species – compartmental segregation, driven by preferential interactions between chromatin states, and loop extrusion, mediated by SMC complexes. These mechanisms provide a tractable and biologically interpretable framework for modeling chromosome structure and dynamics. We note, however, that additional mechanisms—such as transient looping, transcription-mediated interactions, or other non-equilibrium effects—were not explicitly considered and could explain discrepancies between the simulations and the experimental data.

## Conclusion

In this study, we used coarse-grained polymer simulations to systematically evaluate mechanistic models of chromatin folding and reconcile structural and dynamical properties of a *Drosophila* chromosome. Our results showed that neither an ideal chain nor loop extrusion alone could simultaneously reproduce all experimental observables. In contrast, a block copolymer model incorporating experimentally derived genome compartments, particularly near the coil–globule phase transition, significantly improved agreement with experimental data. Adding loop extrusion to the compartment model did not improve significantly the overall agreement across observables.

We drew on recent developments in polymer physics to present evidence supporting that *Drosophila* chromosomes organization and dynamics are consistent with a near-critical folding regime, enabling rapid spatial exploration and frequent long-range enhancer–promoter interactions. Future work should further investigate whether similar near-critical folding regimes operate in mammalian systems and how additional factors—such as boundary elements, epigenetic modifications, and transcriptional activity—modulate chromatin structure and dynamics across scales.

## Materials and Methods

### Computing observables from the experiments

Five key observables from experimental data were used in this study to fit or compare with simulated data. Three of these, i.e., the end-to-end distance R(*s*), the single-locus mean-squared displacement MSD_1_(*t*), and the two-locus mean-squared displacement MSD_2_(*t*), were computed from the live-cell imaging trajectory data of Brückner & Chen et al. [67] using the code originally published with the article [68]. The relaxation time *τ* (*s*) was provided to us by the authors of [42] upon request. Because the live-cell imaging data were obtained from genetically modified embryos containing a synthetic enhancer-promoter system and *homie* elements, we restricted our analysis to trajectories corresponding to open configurations without *homie*-*homie* looping (O_off_ state in the original study). This filtering removes long-lived contacts associated with engineered interactions and allows us to approximate baseline chromatin dynamics that are more directly comparable to polymer models without locus-specific interactions. We verified that this filtering does not qualitatively affect short-time dynamics but reduces the contribution of persistent contacts at large genomic separations (see relevant section below).

The last key observable – the contact probability scaling P_c_(*s*) – was computed using the `expected_cis` function of the *cooltools* package [69] from the experimental Hi-C map (at 16 kb resolution) obtained from another study [52] performed in the same developmental stage of the *Drosophila* embryo as Brückner et al.

### Computing observables from the simulations

From the simulation trajectories comprising the coordinates of at least 7 × 10^4^ polymer conformations sampled over time, R(*s*), MSD_1_(*t*) and MSD_2_(*t*) were calculated for *s* = 58, 82, 88, 149, 195, 595, 3327 kb using Eq. 1, Eq. 2, and Eq. 3 respectively.

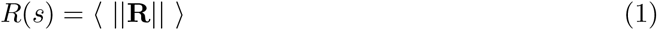

where **R** is the distance vector between the two loci.

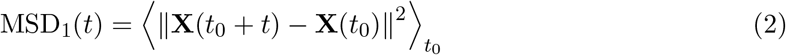

where **X** is the position vector of a locus, *t*_0_ is the initial time, and *t* is the lag time.

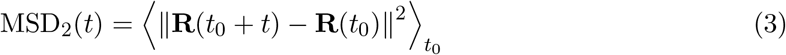

The effect of the localization error of 180 nm in [42, 70] was tested on R(*s*) computed from various simulation trajectories to ensure that this factor could not affect the fits significantly. Indeed, the effects of localization error were negligible (Fig. S5).

The relaxation time *τ* (*s*) was measured from the simulations by computing the time taken for the autocorrelation of the **R** for each *s* (4) to decay to a specific value 5. The scaling exponent was then estimated by taking the slope of *τ* (*s*) plotted on a logarithmic scale.

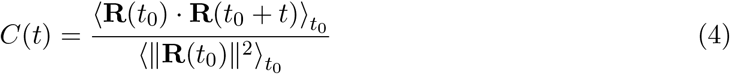

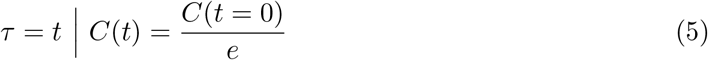

The Hi-C map for each simulation was generated using the `contactmaps.binnedContactMap` function of the *polychrom* package [71], which bins monomer-resolution contacts to 16 kb resolution based on the input contact threshold. The contact map was then normalized using Sequential Component Normalization [72] and used to compute P_c_(*s*).

The value of the contact threshold *k*, used to define contacts in simulated Hi-C maps, can significantly affect the resulting agreement with experimental data. As illustrated in Fig. S6A, different values of *k* lead to noticeable changes in the strength and sharpness of compartmentalization patterns. To assess the robustness of our results, we systematically evaluated the similarity between simulated and experimental Hi-C maps by computing Pearson and Spearman’s correlation coefficients over a range of *k* values (Fig. S6B, C). The highest map-to-map correlations were obtained for *ε* ≈ 0.270–0.285 with *k* = 250 − 300 nm, with a slightly better correlation at *ε* = 0.275 (Pearson *r* = 0.43 for *k* = 250 nm). While the visual agreement between simulated and experimental maps may still appear imperfect, we emphasize that in many previous studies different color scales are used for simulated and experimental maps, which can artificially enhance visual similarity; here, we deliberately use the same color scale for both, providing a more stringent and transparent comparison (Fig. 3D).

Lastly, to compute mean first encounter times (MFETs), several pairs of monomers were selected for each *s* = 50, 100, 250, 500, 1000, 2000 kb along the polymer chain. For the compartment-based block copolymer simulations, each monomer pair was categorized as C0–C0, C1–C1, or C0–C1 based on the compartment into which they fell. Then, for each pair in each simulation, the first encounter time was calculated using 6, starting from an initial time point in which *R >* 100 nm. After binning these values based on *s* and compartment combination, the mean was calculated.

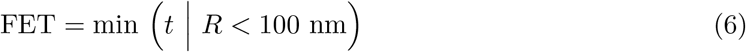

where *R* is the physical distance between the two loci.

The comparison of MFETs from the block copolymer simulations with the reference Rouse model with excluded volume (MFET_ref_) was done by computing the measure given in 7.

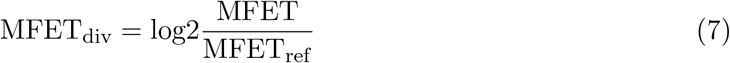

### Fitting observables between simulations and experiments

To fit R(*s*), the curve from experimental data was translated along the y-axis (i.e., the axis corresponding to physical distance) on a logarithmic scale (Eq. 8, equivalent to scaling distances by a numerical factor) until the deviation *σ*_*R*_ between the experimental and the simulated curves reached a global minimum. No translation was necessary along the x-axis because the genomic distance *s* was shared by both data sets. The L-BFGS-B optimization algorithm [73, 74] was used for this fit. This fitting procedure was also used for *τ* (*s*).

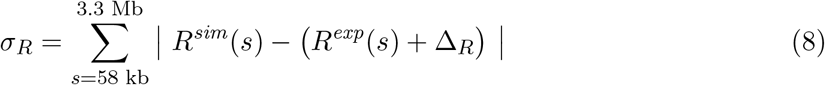

This R(*s*) fit was then used to convert simulation spatial units s.u. (i.e., bead-bead bond length) into real spatial units (i.e., nm) using Eq. 9).

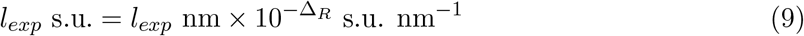

This correspondence allowed us to convert the contact threshold of 250 nm into simulation spatial units, which was then used to compute Hi-C maps. For example, depending on the simulation case, a value of 5 s.u. varied roughly between 132 and 308 nm based on Δ_*R*_ computed from the corresponding R(*s*) fits. In the case of the best-fit for the block copolymer, 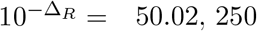 nm was estimated to be ≈ 5 s.u..

Fitting MSD_2_(*t*) required a more elaborate procedure, as the simulated and experimental datasets contained different numbers of points. Each simulated MSD_2_(*t*) curve was first fit to a fifth-degree polynomial ℱ _*s*_ on a logarithmic scale to interpolate the data and enable functional evaluation. Then, using the same optimization algorithm, the experimental data points for each genomic separation *s* were translated along both space and time axes in log–log space—equivalent to fitting both squared displacement and time simultaneously—until the deviation 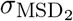 was minimized (Eq. 10).

This procedure was also used for MSD_1_(*t*), with the sole difference being that the initial polynomial fitting used a first-degree instead of a fifth-degree polynomial.

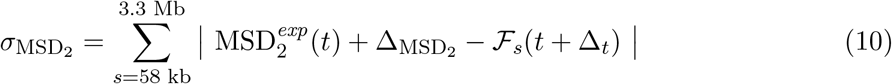

As with spatial units, real time units (in seconds, s) can be converted to simulation time units (timestep of the Langevin integrator, t.u.) and vice versa using the temporal scaling factor Δ_*t*_ obtained from the fitting procedure (Eq. 11). For the best-fit case of loop extrusion, we obtained 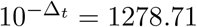, implying that 1 s = 1278.71 t.u.

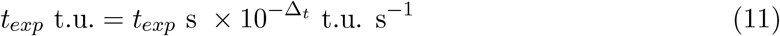

To estimate the deviation 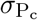 between the simulated and the experimental Hi-C data, we computed the sum of absolute differences between the two curves for equally spaced *s* values on a logarithmic scale. The correlation (both Spearman’s and Pearson) was calculated between the two corresponding observed-over-expected maps as described by Eq. 12.

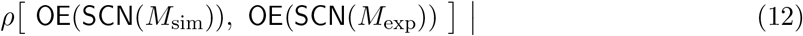

where *M*_*x*_ denotes a Hi-C map, SCN is the normalization function [72], and OE is the observed-over-expected function. The latter involves calculating, for each value on the Hi-C map, the log ratio of the expected value estimated based on its corresponding *s*.

To compute a global minimum for fits to all observables (as performed for the model combining loop extrusion and compartments), deviations for each observable were first normalized, and then combined into a single metric. Specifically, the global deviation was defined as 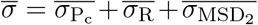, where each 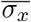 represents the Z-score normalized deviation for observable *x*, computed across all parameter combinations for that simulation case.

### Modeling and simulations

The 4.2 Mb stretch of chromosome 2R (chr2R:6,457,453–10,658,620), encompassing all enhancer–promoter segments, was modeled as a fully flexible homopolymer bead-spring chain—i.e., an *ideal chain*—with a total of 4208 beads, each representing 1 kb of genomic distance. For simplicity, the *MS2* and *parS* insertions used in the experiment were modeled as point insertions along the polymer.

Based on data from [42], promoter loci were assigned to beads 3563, 3439, 3593, 3372, 3695, 2926, and 193. The enhancer was positioned at bead 3505 for downstream promoter loci and at bead 3521 for upstream ones, preserving the experimental enhancer–promoter genomic separations of 58, 82, 88, 149, 190, 595, and 3327 kb.

The conformational dynamics of this ideal chain are described by the Rouse model [54, 75]. In all models except the Rouse model, excluded volume interactions were introduced using a parabolic repulsive potential, as defined in Eq. 13.

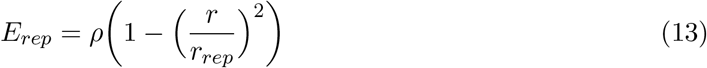

where the maximum height of the potential was *ρ* = 5 kT and its width was *r*_*rep*_ = 1

For all models, the thermal fluctuations of the polymer chain over time were simulated under overdamped Langevin dynamics [76] using the OpenMM-based *polychrom* package [71, 77]. The simulations of the Rouse model with and without excluded volume were first equilibrated for 10^7^ t.u., then the chain dynamics were recorded for 10^9^ t.u., with one conformation being sampled every 10^3^ t.u.. For all other simulations, these values were 8 × 10^6^ t.u., at least 5.6 × 10^8^ t.u. and 8 × 10^2^ t.u. respectively. For the simulations of the Rouse model, the only parameters used were the equilibrium bead-bead harmonic bond length (1 s.u.) and its standard deviation (≈ 0.28 s.u.); the parameters for the other models are discussed in the sections below.

#### Loop extrusion

A common two-step framework was used to simulate loop extrusion on the chromosome [13, 58]. For each case, first, a stochastic simulation of LEF dynamics was performed on a onedimensional lattice of the same length as the polymer. LEFs could stochastically load onto and unload from the lattice. They were allowed to slide across the lattice, and while doing so, could cross one another or collide and stall. The loading, unloading, sliding, and crossing of LEFs were dictated by a set of loop extrusion parameters (Table 1). No insulators or boundary elements were introduced in any of the simulation cases, and therefore, the LEFs moved freely unless they were stalled by another LEF. During the simulation, the positions of the LEFs on the lattice were recorded over time to keep track of the loops formed and dissolved. Next, during the 3D Langevin dynamics simulation of the polymer, beads corresponding to the LEF positions from the lattice-based simulation trajectory were brought together and anchored by harmonic bonds to mimic the bridging action of LEFs, thus forming loops in the chain. By constantly moving the loop anchors to beads that matched the positions on the lattice-based trajectory, dynamic loop extrusion was simulated on the polymer.

**Table 1.** Parameters of loop extrusion for stochastic simulations on the lattice.

| Parameter | Value |
| --- | --- |
| Lattice length | 4208 |
| LEF unloading probability | 0.01 |
| LEF loading probability | 0.08 to 0.8, stepping 0.08 |
| LEF sliding probability | 0.5 |
| LEF crossing probability | 0.5 |
| LEF anchor length | 1.25 s.u. |
| Time units for LEF probabilities | 800 t.u. |

The stochastic model was implemented such that the ratio between the probabilities of LEF loading *p*_*L*_ and unloading *p*_*U*_ determined the steady-state mean occupancy of LEFs ⟨Ω⟩ on the lattice (Eq. 14). The steady state was dependent on the size of the lattice and the sliding probability of the LEFs *p*_*S*_, and was achieved for time *t >>* 10^4^ t.u. Therefore, to vary ⟨Ω⟩, *p*_*L*_ was varied while keeping the sliding probability *p*_*S*_ and *p*_*U*_ constant. *p*_*S*_ = 0.5 and *p*_*U*_ = 0.01 were assigned to attain a mean LEF processivity 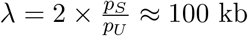.

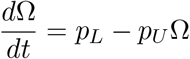

Steady state achieved when 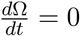, therefore

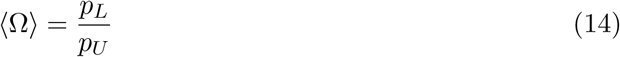

From Eq. 11 and the average time taken for a LEF to slide 1 kb (800 t.u., Table 1), LEF parameters were estimated in real units. Depending on the simulation case, 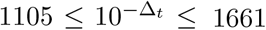 t.u. s^−1^. Taking the best-fit case for dynamics (⟨Ω⟩ = 0.95 LEFs per 100 kb), we find

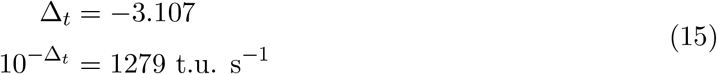

Using this conversion factor, we estimated the values for mean LEF velocity as v_L_EF ≈ kb s^−1^ (Eq. 16) and mean LEF residence time as *τ*_LEF_ ≈ 63 s (Eq. 17).

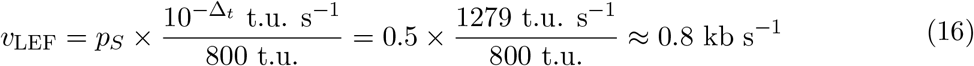

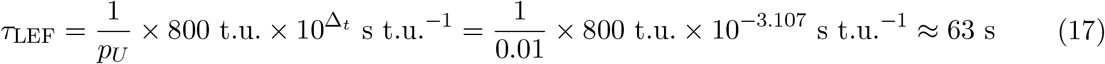

#### Compartmental segregation

The compartments of the chromosomal segment were predicted by extracting the first eigenvector of the genomic distance-normalized experimental Hi-C map at 16 kb resolution [52]. A large region of the chromosome harboring the loci of interest but excluding the centromere (chr2R:6,464,000–25,286,936) was used for the computation. Each 16 kb segment of this stretch was associated with one compartment (named C0 or C1). Each bead in the bead-spring chain with excluded volume was assigned a type based on the compartment predicted, resulting in blocks of alternating C0 and C1 beads. Attractive interactions were introduced only between beads of the same type during Langevin dynamics simulations. The strength of attraction between the beads was controlled by the parameter *ε*, which corresponds to the depth of the attractive well in a custom parabolic potential (Eq. 18). This potential was devised to induce strong attraction between beads at a short range.

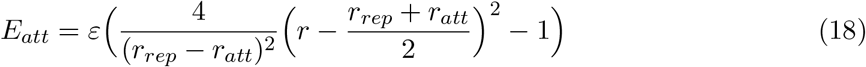

where 0.18 ≤ *ε* ≤ 0.32 and the width as measured from *r* = 0 was *r*_*att*_ = 1.8

#### Test simulation with *homie* elements

To test the effect of *homie* elements and the subsequent filtering of looped trajectories on the computation of experimental observables, a simple test simulation was performed. A Rouse polymer with a pair of *homie* elements (single monomers) placed 595 kb apart was simulated, using the same attractive potential as the block copolymer (Fig. 3A, with *r*_*att*_ = 3 s.u. and *ε* = 7) to introduce looping upon proximity. Other monomers in the polymer did not have any specific interactions. The simulation trajectory was filtered using *k* = 2.5 s.u. to remove *homie*-*homie* loops (Fig. S7A), splitting the trajectory into smaller ones (Fig. S7B), analogous to the O_off_ state in [42]. The distribution of R(*s*), and the MSD_1_(*t*) and MSD_2_(*t*) curves in Fig. S7C–E, respectively, showed negligible differences between the Rouse model and the *homie* simulation after filtering.

### Testing criticality-driven dynamics

#### Measurement of polymer fluctuations

The fluctuations in polymer size over the simulation trajectory were measured by computing the standard deviation of the squared radius of gyration (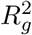 , Eq. 19) of the polymer conformations in the ensemble, and dividing it by the mean 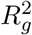. To compute this measure for a compartment block, the same computation is applied to only that segment of the polymer.

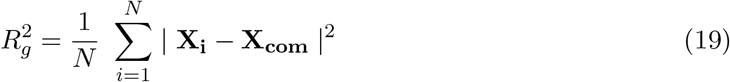

where *N* is the number of monomers, **X**_**i**_ is the coordinate vector, and **X**_**com**_ is the center of mass of the polymer.

#### Computation of power spectral density and *α*

The power spectral density (PSD) of an ensemble of polymer conformations and its slope *α* were calculated using code available from [78]. First, the squared discrete cosine transform was applied to the x-, y-, and z-axis spectra (i.e., projections) of each conformation in the ensemble, which was then averaged to yield the PSD. Plotting the PSD as a function of its polymer mode on a logarithmic scale allowed the extraction of *α*, which is the slope of the first three spectral modes. *α* is an indicator of the polymer ensemble’s state or position within the coil–globule transition [63], starting from *α* ≈ −2.2 for a swollen coil and *α* ≈ −2 for an ideal chain, increasing to *α* ≈ 0 for a globule.

## Data and code availability

The simulation-launcher code is available at https://github.com/ggautham07/polychrom-sim-launchers, and the analysis code is available at https://github.com/ggautham07/eve-project-analysis-2026.

## Acknowledgments

We are grateful to Maria Barbi, Timothy Földes, Thomas Gregor, David Llères, and Marie Schaeffer for critical reading of the manuscript. We thank Alessandro Barducci, Maria Barbi, Timothy Földes, Andrea Parmeggiani, Daniel Jost, and Antoine Coulon for valuable discussions. G.G was funded by the Doctoral School for Chemical and Biological Sciences for Health (EDCBS2) at the University of Montpellier. We acknowledge the PCIA computing platform at the National Museum of Natural History, Paris. This project was funded by the European Union’s Horizon 2020 Research and Innovation Program (Grant ID 724429, M.N.) and the French National Research Agency (ANR-23-CE12-0023-01, M.N.).

## Supplementary Figures

**Figure S1:**
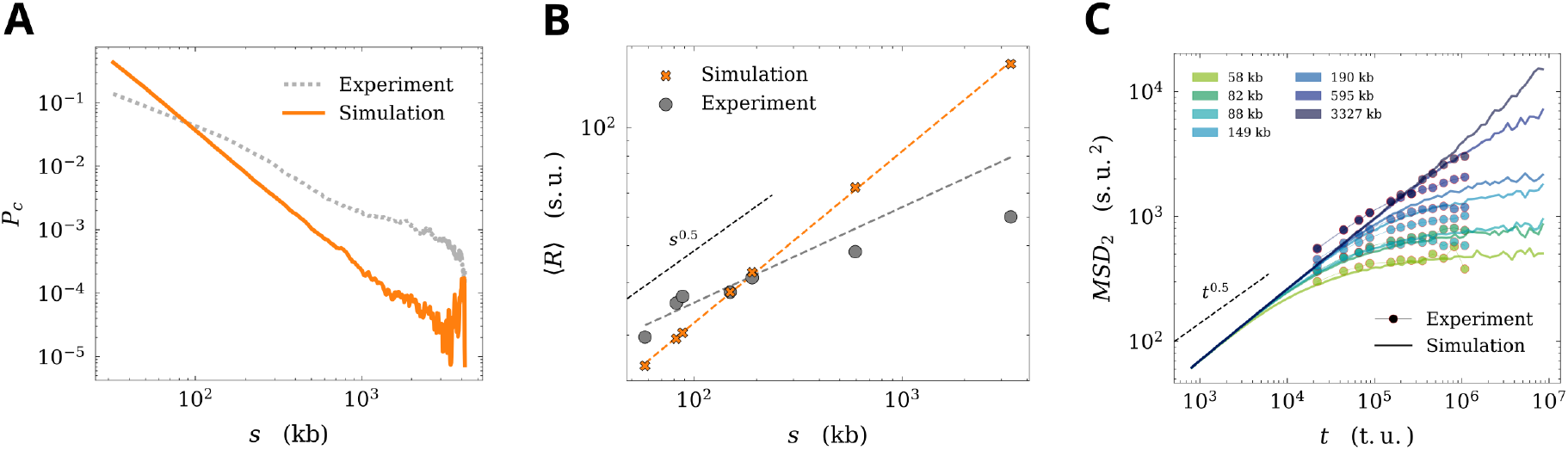
Fits to the experimental data for the Rouse model with excluded volume (i.e., a swollen coil) simulation. **A** – P_c_(*s*), **B** – R(*s*) (*β* = 0.61), **C** – MSD_2_(*t*).

**Figure S2:**
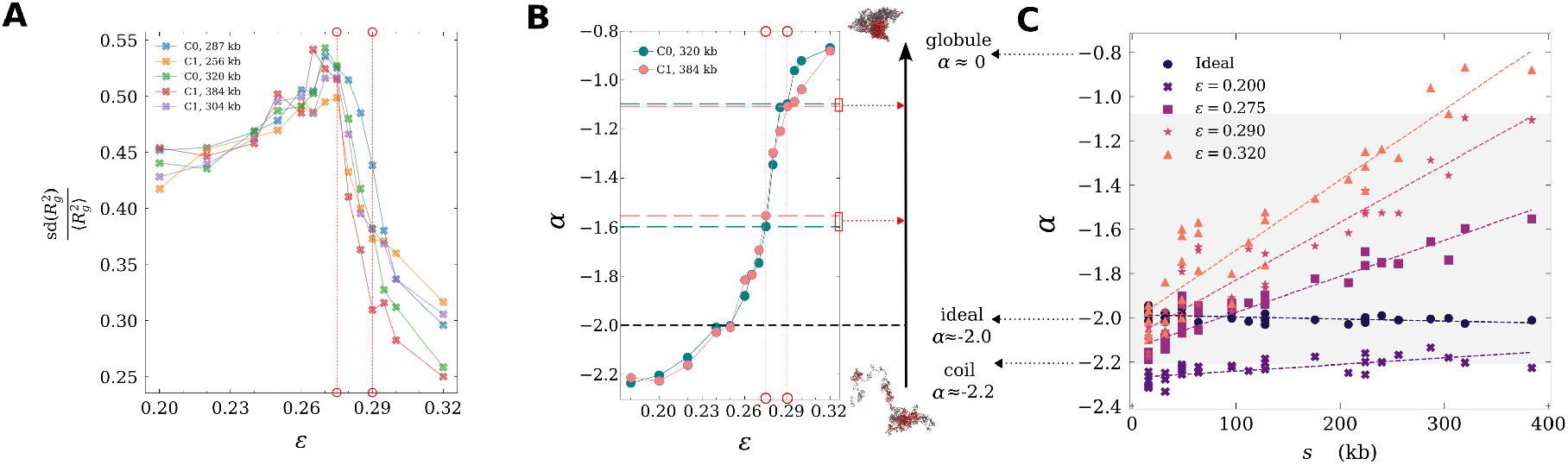
Evidence for near-critical polymer behavior for compartment domains, **A** – fluctuations in the mean-normalized standard deviation, **B** – slope of the power spectral density *α* for the two largest compartment domains, **C** – Scatter plot of *α* values for all compartment domains as a function of their genomic size.

**Figure S3:**
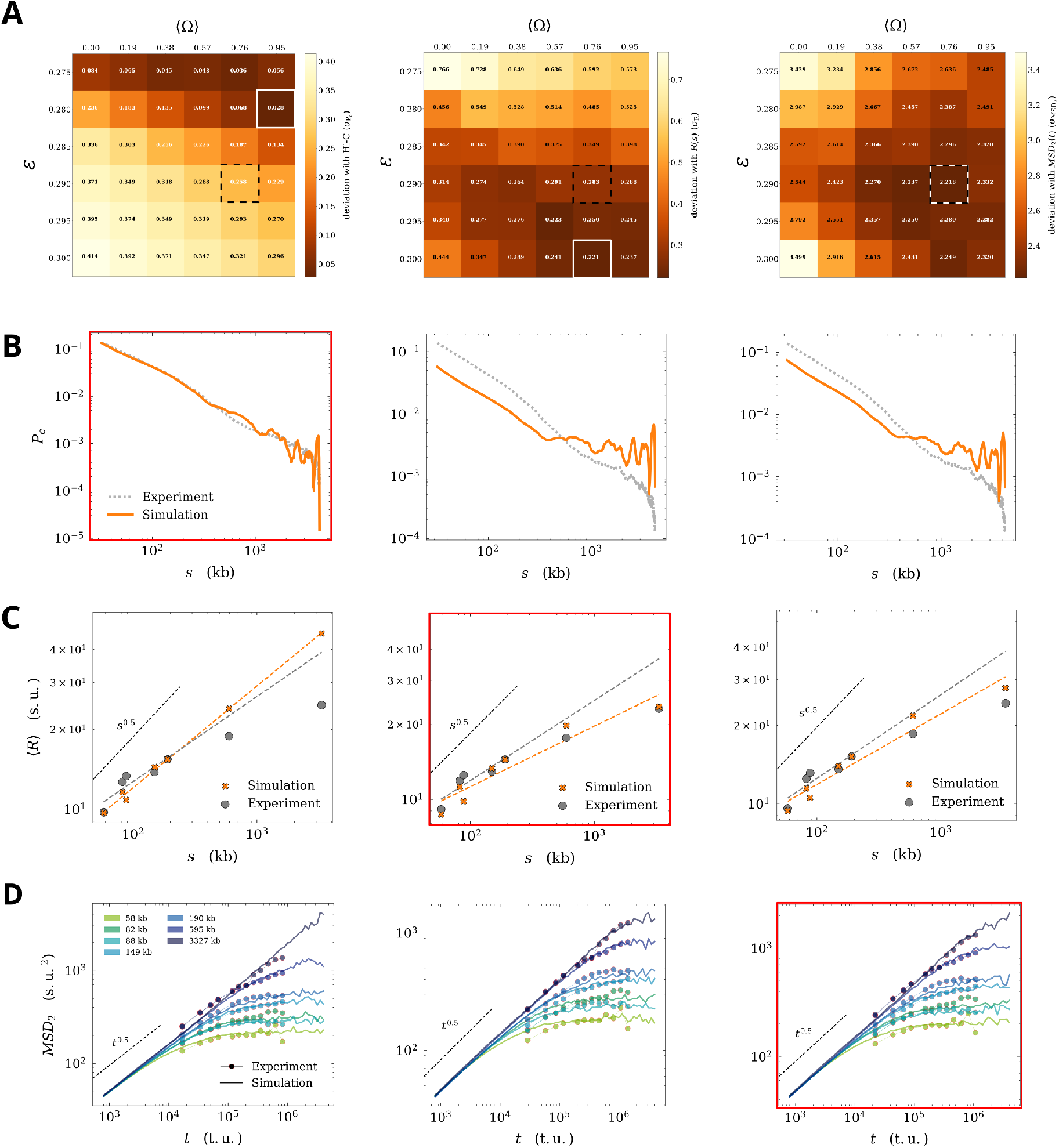
**A** – Grids showing 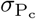 (left), *σ*_R_ (center), and 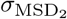 (right) as a function of parameter pairs of *ε* and ⟨Ω⟩ for the compartment-based block copolymer with loop extrusion. White boxes indicate the best fit corresponding to each observable. Dashed black boxes highlight the global minimum computed in Fig. 5. **B** – Fits for P_c_(*s*), R(*s*), and MSD_2_(*t*) for the simulation with the lowest 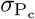. **C** and **D** – Same for *σ*_R_ and 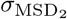 respectively. The fit corresponding to the minimum of each observable is marked in red along each column.

**Figure S4:**
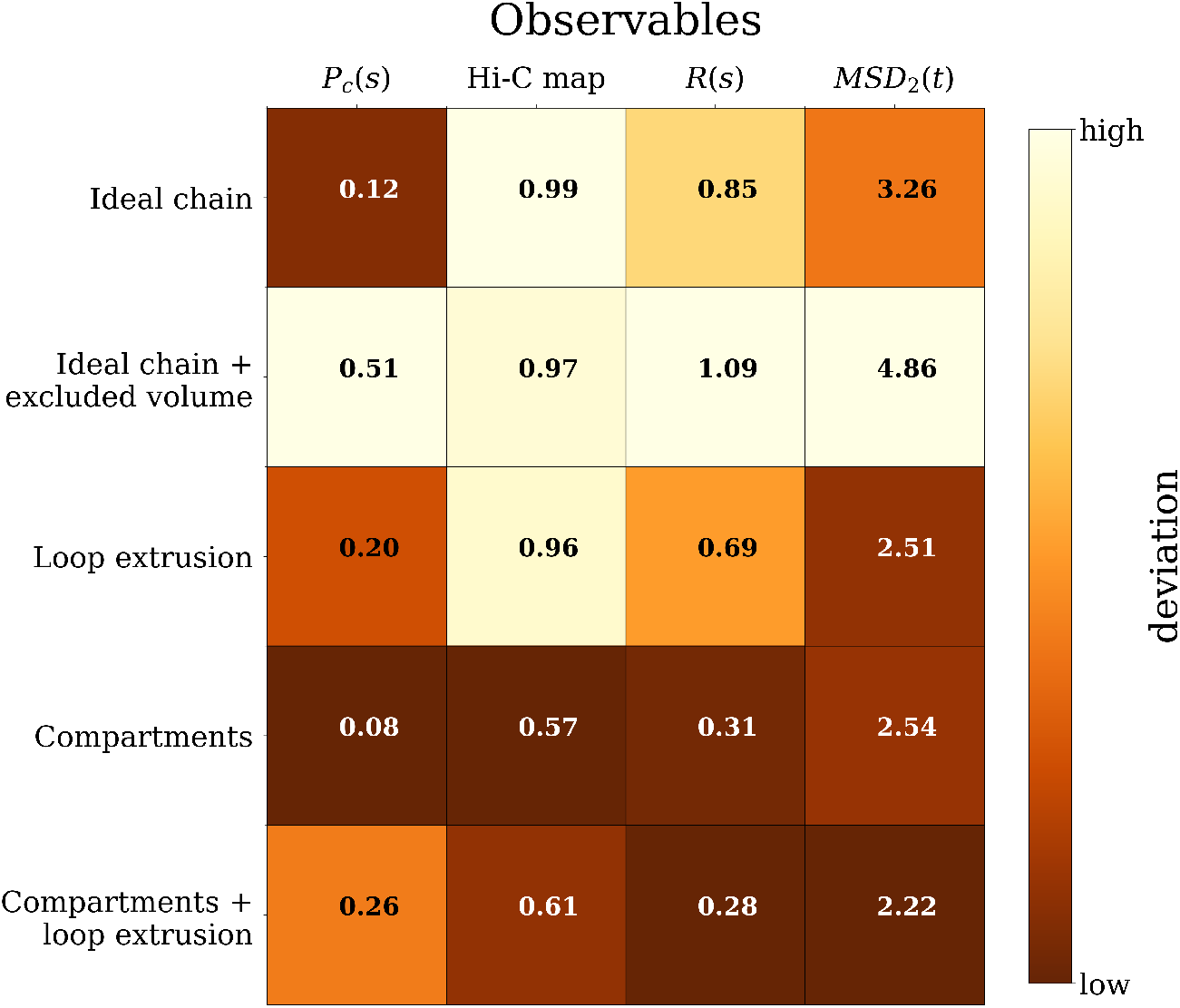
Summary grid with deviations for all observables: P_c_(*s*), 1 - Pearson with Hi-C map, R(*s*) (absolute deviation in logarithmic scale), and MSD_2_(*t*) (same) across all explored models

**Figure S5:**
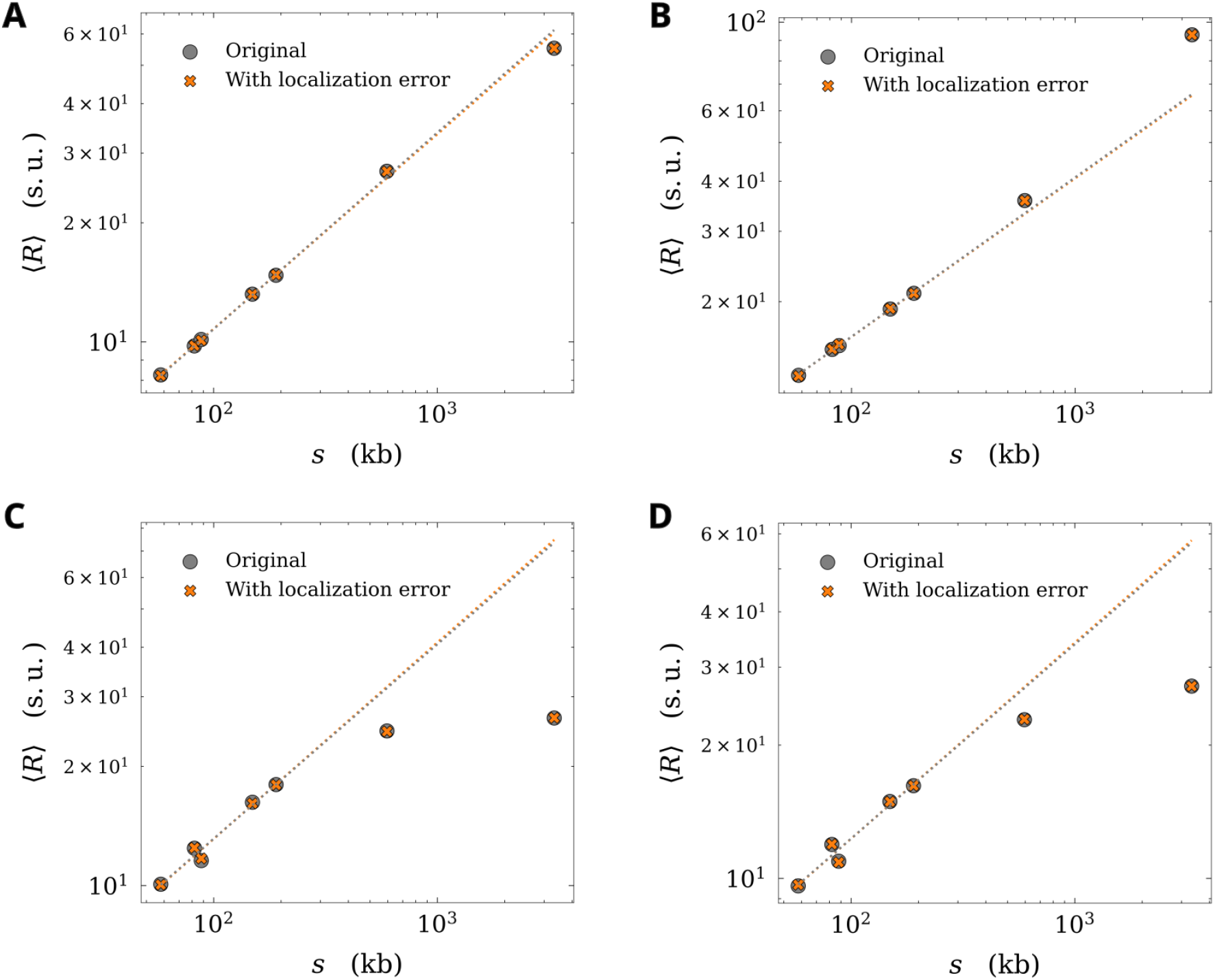
Effect of localization error on ⟨*R*⟩ is negligible for all simulations, **A** – Rouse model, **B** – Best-fit loop extrusion, **C** – Best-fit compartment-based block copolymer, **D** – Global best-fit for loop extrusion on compartment-based block copolymer

**Figure S6:**
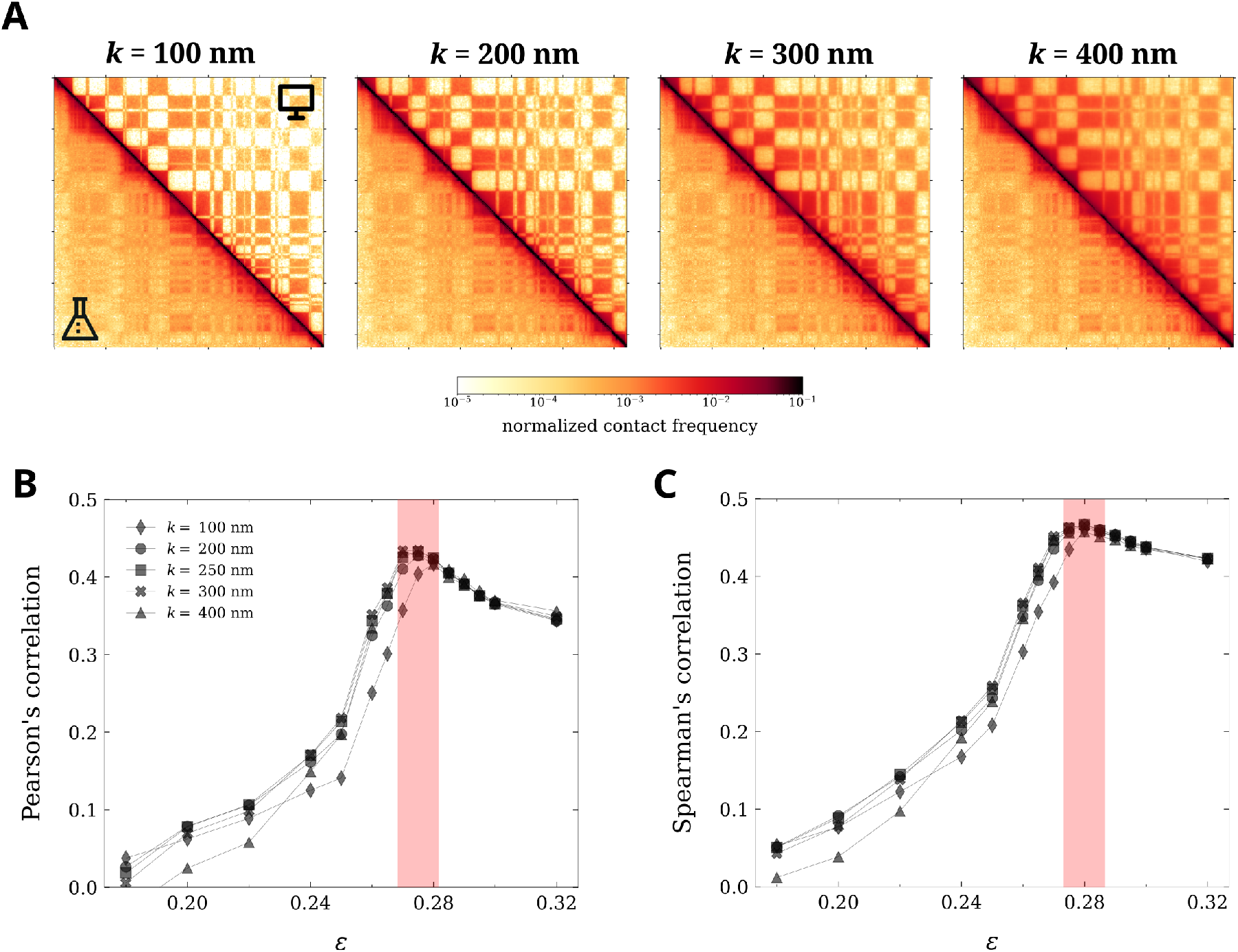
Choosing a suitable cut-off from comparisons with Hi-C data for the block copolymer simulations. **A** – Hi-C map computed from the simulation with *ε* = 0.275 using an increasing contact threshold *k* in nm. **B** – Pearson correlation with the Hi-C map computed using different *k* values as a function of *ε*. **C** – same computation with Spearman’s correlation.

**Figure S7:**
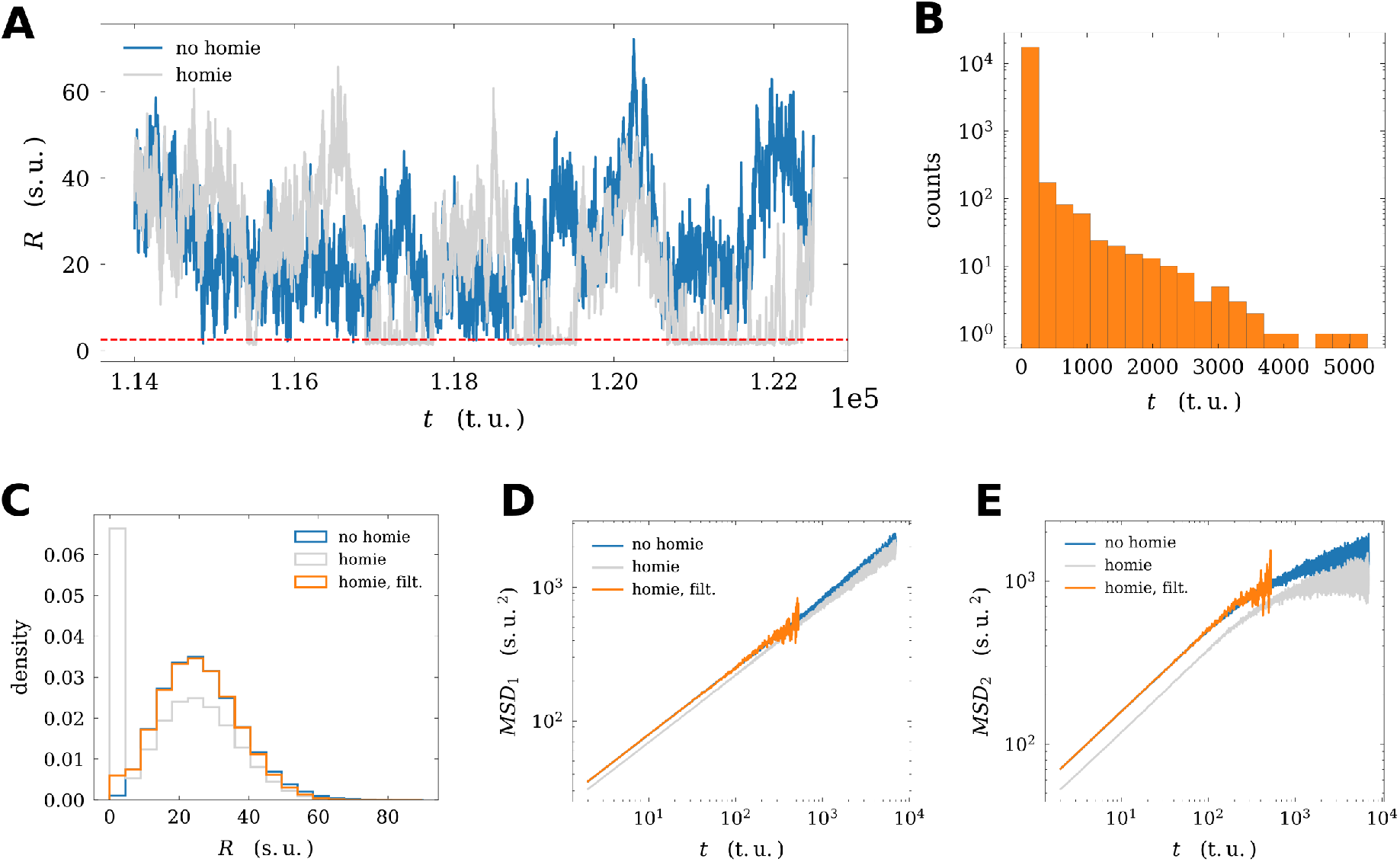
Effect of filtering *homie*-looped configurations. **A** – sample of simulation trajectory showing differences between the Rouse model (“no homie”) and the Rouse model with one pair of attracting *homie* elements placed 595 kb apart (“homie”). **B** – distribution of the lengths of new trajectories after filtering looped trajectories (remove all *R <* 2.5 s.u., called “homie, filt.”) from the original “homie” simulation trajectory. **C** – distribution of *R* for 595 kb from simulations for the three cases: “no homie”, “homie”, and “homie, filt.” **D** – MSD_1_(*t*) computed from each case. **E** – MSD_2_(*t*) computed for each case.

## Notes

### Competing Interest Statement

The authors have declared no competing interest.

### Summary of Updates

This version of the manuscript has been revised to improve the evidence of a near crtitical behavior

